# Model checking to assess T-helper cell plasticity

**DOI:** 10.1101/012641

**Authors:** Wassim Abou-Jaoudé, Pedro T. Monteiro, Aurélien Naldi, Maximilien Grandclaudon, Vassili Soumelis, Claudine Chaouiya, Denis Thieffry

**Author notes:** these authors contributed equally to this work (Wassim Abou-Jaoudé, Pedro T. Monteiro).

## Abstract

Computational modeling constitutes a crucial step towards the functional understanding of complex cellular networks. In particular, logical modeling has proven suitable for the dynamical analysis of large signaling and transcriptional regulatory networks. In this context, signaling input components are generally meant to convey external stimuli, or environmental cues. In response to such external signals, cells acquire specific gene expression patterns modeled in terms of attractors (*e.g.* stable states). The capacity for cells to alter or reprogram their differentiated states upon changes in environmental conditions is referred to as cell plasticity.

In this article, we present a multivalued logical framework along with computational methods recently developed to efficiently analyze large models. We mainly focus on a symbolic model checking approach to investigate switches between attractors subsequent to changes of input conditions.

As a case study, we consider the cellular network regulating the differentiation of T-helper cells, which orchestrate many physiological and pathological immune responses. To account for novel cellular subtypes, we present an extended version of a published model of T-helper cell differentiation. We then use symbolic model checking to analyze reachability properties between T-helper subtypes upon changes of environmental cues. This allows for the construction of a synthetic view of T-helper cell plasticity in terms of a graph connecting subtypes with arcs labeled by input conditions. Finally, we explore novel strategies enabling specific T-helper cell polarizing or reprograming events.

## 1 Introduction

Cellular signaling pathways and regulatory circuits are progressively deciphered, with a recent acceleration allowed by the development of powerful high-throughput experimental approaches. Computational modeling constitutes a crucial step towards the functional understanding of the resulting intertwined networks. Different formalisms have been commonly used to model complex biological networks, with different levels of abstraction (Albert et al., 2013; de Jong, 2002; Karlebach and Shamir, 2008; Samaga and Klamt, 2013). Among these formalisms, the discrete, logical approach is particularly useful to model biological systems for which detailed kinetic data are lacking, which is often the case (Bornholdt, 2008; Naldi et al., 2014; Wang et al., 2012). Moreover, logical modeling allows the consideration and the dynamical analysis of comprehensive signaling/regulatory networks. Here, we rely on the multivalued formalism initially introduced by R. Thomas and collaborators (Thomas and D’Ari, 1990).

Following R. Thomas, we model networks in terms of a *logical regulatory graph*, where nodes represent regulatory components, while edges denote regulatory interactions (activations or inhibitions). Each component is associated with a discrete variable denoting its (current) functional level of activity. In addition, a *logical rule* (or *logical function)* describes the evolution of this level, depending on the values of the regulators of the component. The regulatory graph together with the logical rules enable the computation of the dynamical behavior of the model, which is usually represented in terms of a *State Transition Graph* (STG), where each node represents a *state* of the system (*i.e.*, a vector listing the values of all the variables), while arcs represent enabled *state transitions*. The terminal strongly connected components of a STG denote the attractors of the underlying network, *i.e.* capture its asymptotic behavior in terms of stable states or (potentially complex) dynamical cycles. Consequently, the identification of these attractors and the evaluation of their reachability from given initial condition(s) are paramount to understand network behaviors. However, as the number of states increases exponentially with the number of components, advanced computational methods are needed to analyze the dynamics of discrete models. In this respect, several strategies have been developed to efficiently assess dynamical properties of comprehensive logical models.

Here, we focus on the analysis of networks encompassing input components that embody external signals, instructing intertwined signaling pathways with feedback regulations. Each (fixed) combination of input values (*i.e.* environmental cues) defines a specific region of the state space where the dynamics and its associated attractors are confined. In the case of models of networks controlling cell differentiation, attractors correspond to differentiated patterns of gene expression (or protein activity). We call *differentiated states* these attractors, which are generally stable states (see *e.g.* Naldi et al. (2010)), but can also be complex attractors denoting homeostasis or oscillatory behavior (see *e.g.* Bonzanni et al. (2013)). It is of particular interest to assess how input value changes affect differentiated states, sometimes resulting in functional reprograming. The capacity of cells to change their asymptotic behaviors depending on environmental cues is referred to as *cell plasticity* (see *e.g.* O’Shea and Paul (2010)). In this manuscript, we present a methodology to assess cell plasticity, relying on the logical formalism assets and recent computational methods, including model checking techniques.

Model checking is a computer science technique for the verification of large discrete dynamical systems (Clarke et al., 1999). It has been recently applied to the analysis of biological networks (Arellano et al., 2011; Batt et al., 2005; Brim et al., 2013; Chabrier and Fages, 2003; Schwarick and Heiner, 2009). Properties are formalized in terms of temporal logic statements, and the verification process explores (restricted) regions of the state space, in order to check the truthfulness of the properties. Here, we consider a further improvement that consists in defining input values as labels of the transitions in STGs, thereby reducing the number of states. This allows to efficiently assess input conditions when verifying, for example, reachability properties between differentiated states. For this, we use a specific symbolic model checker called *NuSMV-ARCTL*, along with temporal logical semantics enabling the specification of properties with restrictions on the input valuations (Lomuscio et al., 2007).

We consider the case of T-helper cell differentiation to demonstrate the assets of the logical framework and the power of model checking to elucidate how cells respond to environmental stimuli. More precisely, we model the cellular network controlling the differentiation of CD4 T-helper (Th) cells, which regulate many physiological and pathological immune responses. Upon activation by antigen presenting cells (APC), naive Th cells polarize into distinct Th subtypes expressing different sets of cytokines, tailoring appropriate immune responses to the invading pathogen. Recent experimental data highlight the ability of Th subsets to alter and even reprogram their phenotypes, according to environmental cues (Nakayamada et al., 2012). These observations challenge the classical linear view of Th differentiation into distinct separated lineages, raising fundamental questions regarding the mechanisms underlying Th differentiation and plasticity.

In order to get insights into the dynamical behavior of T-helper cell differentiation, several models describing the regulatory network controlling Th commitment have been proposed, relying on quantitative modeling approaches (Mendoza and Pardo, 2010; van den Ham and de Boer, 2008, 2012) or using discrete qualitative frameworks (Martinez-Sosa and Mendoza, 2013; Mendoza, 2006; Naldi et al., 2010). Here, the logical model of Th cell differentiation of Naldi et al. (2010) is extended to cover several novel Th subtypes. Focusing on Th polarization and reprograming events, we show how biologically relevant properties can be formalized and tested using model checking. More precisely, we compute all reprograming events between Th subsets under specific documented polarizing cytokine environments, providing a global and synthetic representation of Th plasticity in response to these environmental cues. This analysis leads to the prediction of Th subtypes conversions, which would need to be assessed experimentally. Finally, we delineate several strategies for Th subtype reprograming, as well as for naive Th cell polarization towards a novel hybrid Th subtype (predicted by our model).

This manuscript is organized as follows. Section 2 briefly reviews the basics of the logical modeling framework, including model definition and an overview of computational methods to analyze dynamical properties. We also introduce the use of model checking to enhance the analysis of logical models, in particular when these include input components. This methodology is then applied to a logical model for Th differentiation in Section 3, which includes a presentation of the resulting biological insights. Section 4 concludes the manuscript with a discussion and some prospects.

## 2 Materials and Methods

In this section, we introduce the logical framework, presenting the rationale underlying model definition. We further describe model modifications accounting for genetic perturbations (*e.g.*, gene knock-out or knock-in) along with a model reduction method. Next, we briefly present computational strategies to efficiently analyze properties of logical models. Finally, we focus on the assets of model checking to enhance the dynamical analysis of large signaling/regulatory logical networks. Figure 1 illustrates the workflow for logical model definition and analysis, on which we rely to address the question of Th cell plasticity. Most methods presented in this section are implemented in GINsim (http://ginsim.org, Chaouiya et al. (2012)).

**Figure 1.**
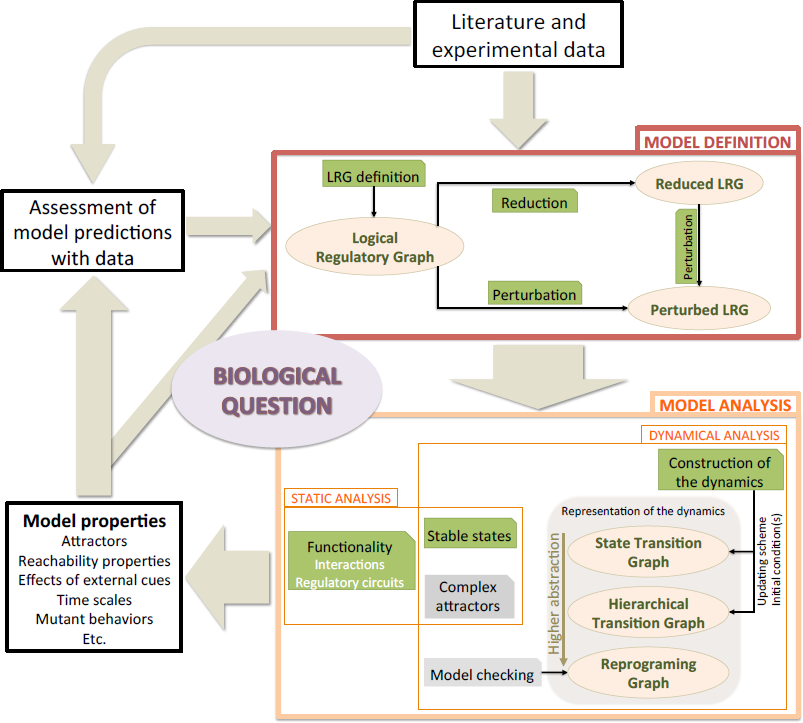
Typical workflow to tackle a central biological function using logical model construction and analysis. A model is defined, relying on literature and experimental data (box *Model Definition).* The model is then analyzed (boxes *Static analysis* and *Dynamical analysis*). The identification of the attractors is performed either by static methods (see Sections 2.2.1 and 2.2.2) or by inspecting the dynamics (see Sections 2.2.3 and 2.3). Dynamics are represented at different levels of abstraction, from the comprehensive state transition graphs to the reprograming graphs. Resulting properties are confronted with biological observations, leading to predictions and/or to model revision. Ellipsoid boxes relate to the different model versions and behavior representations. Green boxes denote methods that are available in GINsim, whereas gray boxes denote analyses performed with other software tools.

### 2.1 Logical Model construction

This subsection shortly introduces the definition of multivalued logical models (for more details and formal definitions, see *e.g.*, Chaouiya et al. (2003); Thomas and D’Ari (1990)).

#### 2.1.1 Logical formalism

A logical model of a regulatory and/or signaling network is defined as a *Logical Regulatory Graph* (LRG), where:

- {*s*_1_, … *s*_*n*_} is the set of nodes, which embody the components of the network; these may correspond to proteins, genes, or phenomenological signals (*e.g.*, the node APC in Figure 2 denotes an antigen presenting cell).
- Each component *s*_*i*_ is associated with a discrete (positive integer) variable, which takes its values in *S*_*i*_ = {0,…, *max*_*i*_}; for simplicity, we denote both the component and its associated variable by *s*_*i*_, embodying the component level of activity or concentration. In general, the maximum level of *s*_*i*_, denoted *max*_*i*_, is set to 1 (*i.e.*, Boolean variable), but it can take higher values to convey qualitatively distinct functional levels.
- Each interaction (*s*_*i*_, *s*_*j*_, *θ*) is defined by its source *s*_*i*_, its target *s*_*j*_ and a threshold *θ*; the interaction is said to be effective when *s*_*i*_ ≥ *θ*; note that *θ* ≤ *max*_*i*_ (the threshold cannot exceed the maximal level of the source).
- The state space of the LRG is given by 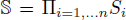; hence a state of the model is a vector s = (*s*_*i*_)_*i*_=_1,…,*n*_.
- The model behavior is specified in terms of *logical rules* (or *logical functions*): the evolution of *s*_*i*_ is defined by 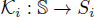 with 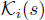 specifying the target value of *s*_*i*_ when the system is in state *s*.

**Figure 2.**
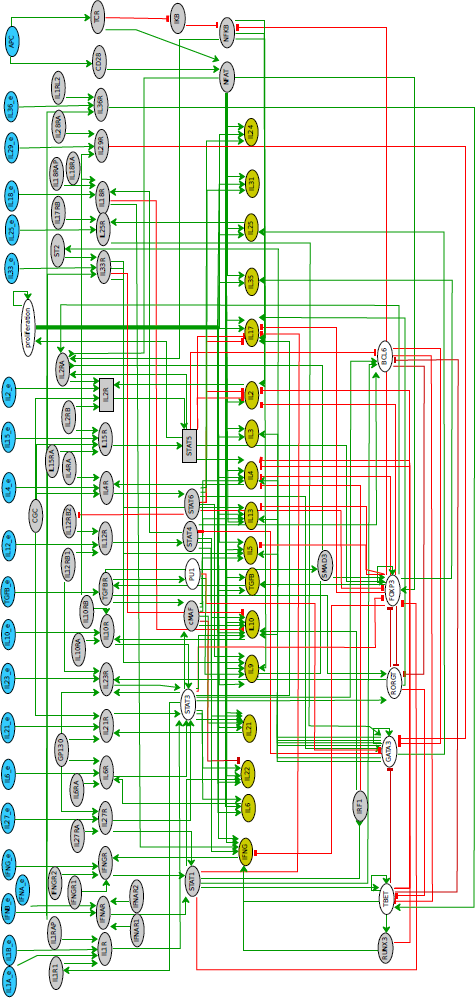
Regulatory graph of Th differentiation logical model. The model encompasses 101 components (among which 21 input nodes) and 221 interactions. The components denoting the inputs are in blue, those denoting the secreted cytokines in orange. Green edges correspond to activations, whereas red blunt ones denote inhibitions. Ellipses denote Boolean components, whereas rectangles denote ternary ones. Grey-out components are those selected for reduction.

The software GINsim provides a graphical interface for the LRG definition, including the components (nodes) and their ranges (maximum values), the interactions (signed arcs) and their thresholds, along with the logical rules (using Boolean expressions or *logical parameters* (Thomas and D’Ari, 1990)).

The behavior of an LRG is classically represented in terms of a *State Transition Graph* (STG), which encompasses the initial model state(s) together with their direct and indirect successors. A transition between two states corresponds to the update of specific components. These updates are dictated by the logical rules. When several components are called to change their values at a given state, these updates are performed according to an updating scheme. The most used updating schemes are the fully synchronous updating (all changes are performed simultaneously, leading to a unique successor), and the fully asynchronous updating (all changes are performed independently, leading to as many successors as the number of updated components). Further details on STG and updating schemes are provided in Section 2.2.3.

In such dynamical models, the asymptotic behavior of the system is captured by the *attractors.* These correspond to the terminal *Strongly Connected Components* (SCC) of the state transition graph. A SCC is defined as a maximal set of mutually reachable states. A SCC is denoted *terminal* when no transition leaves this state set (*i.e.*, once the system enters this set, it is trapped there forever). An attractor is defined by either a single state, which corresponds to a *stable state* denoting a stable pattern of expression often interpreted as a cell differentiation state, or by a larger set of states involved in a dynamical terminal *cycle*, denoting an oscillatory (or homeostatic) behavior. It is therefore important to identify these attractors along with reachability properties (*e.g.*, to determine the attractors reachable from a specific initial state).

#### 2.1.2 Logical modeling of network perturbations

In the logical framework, it is straightforward to define perturbations such as gene knock-out, gene knock-in, or more subtle perturbations (*e.g.*, rendering a component insensitive to the presence of one of its regulators). Modeling such perturbations amounts to specific modifications of the corresponding logical rules. Modifications affecting several components can be easily combined. Given a logical model, one can thus define various perturbations to account for experimental observations or to generate predictions regarding the dynamical role of regulatory components or interactions.

#### 2.1.3 Reduction of logical model

It is often useful to simplify large models by abstracting components, hence diminishing the size of the model state space. In this respect, GINsim implements a reduction method automating the reduction of any component, except those that are self-regulated (Naldi et al., 2011). The computation of a reduced model is performed iteratively: to remove a component, the logical rules of its targets are modified to account for the (indirect) effects of the regulators of this component. This is efficiently done in time polynomial in the numbers of targets (components regulated by the removed one) and regulators of the removed components. In the case of a Boolean model, removing *n* components leads to a reduction of the state space by a factor 2^*n*^.

Obviously, such a reduction may change the dynamics. In fact, it conserves the nature (and number) of the stable states and of the terminal elementary cycles (also called simple cycles, with neither repeated states nor repeated transitions (Berge, 2001)). However, oscillatory components may be split or isolated, and reachability properties only partly conserved. Depending on the type of components that are removed upon reduction, specific dynamical properties are preserved. In Saadatpour et al. (2013), the authors showed that all the attractors of an asynchronous Boolean model are conserved upon reduction of input and pseudo-input components (*i.e.* components with no regulators or regulated by only input and pseudo-input components). Additionally, Naldi et al. (2012) proved that the reduction of output and pseudo-output components not only preserves the attractors, but also their reachability properties (output components regulate no other components, and pseudo-ouput components are those regulating only (pseudo-) output components). In all cases, the existence of a trajectory for a reduced model is maintained for the original model. Hereafter, we take advantage of this reduction method to ease the analysis of our T-helper cell differentiation model (see Section 3).

### 2.2 Model analysis

Means to investigate the dynamical properties of a model can be subdivided into: (1) static analyses, which infer properties without requiring the construction of the STG; and (2) dynamical analyses, which explore proper representations of the dynamics (see Figure 1).

#### 2.2.1 Static analysis — interactions and circuit functionality

The delineation of logical rules for components targeted by several regulators can be relatively tricky. These rules are encoded in GINsim as Multivalued Decision Diagrams, which represent multivalued functions as direct acyclical graphs allowing efficient manipulations (Kam et al., 1998; Naldi et al., 2007).

To help the modeler, GINsim provides a method to check the coherence of the interactions (including their signs) encoded in a regulatory graph with the logical rules associated with its components. Basically, for each interaction (*s*_*i*_, *s*_*j*_, *θ*), GINsim compares the target level of *s*_*j*_ given by its logical function, when (*s*_*i*_, *s*_*j*_, *θ*) is effective (*s*_*i*_ ≥ *θ*) and when it is not (*s*_*i*_ < *θ*), for all combinations of the remaining regulators of *s*_*j*_ (if any). If both target levels are always equal, we say that this interaction is not *functional.* Relying on this comparison, it is also possible to derive the sign of the interaction (activation or inhibition).

*Regulatory circuits* (*i.e.*, elementary cycles in the LRG, also called *feedback loops*) drive non-trivial behaviors such as multi-stability (in the case of positive circuits, involving an even number of negative regulations) or sustained oscillations (negative circuits, involving an odd number of negative regulations) (Thieffry, 2007). Based on the aforementioned method to assess interaction functionality, GINsim enables the delineation of the *functionality context* (if any) of each regulatory circuit (Naldi et al., 2007; Remy and Ruet, 2008). This functionality context is defined as the levels of external regulators that allow each circuit interaction to be functional and thereby affect its target in the circuit. It can be interpreted as the region of the state space where the circuit generates the corresponding dynamical property. This definition enables the identification of the regulatory circuits playing the most important regulatory roles within a complex LRG (see Comet et al. (2013) for further discussion on circuit functionality).

#### 2.2.2 Static analysis – identification of dynamical attractors

Attractors (stable states or terminal cycles) constitute crucial dynamical properties of the model and have thus been the focus of many computational studies. In particular, a SAT-based algorithm was proposed in Dubrova and Teslenko (2011) to compute all the attractors of synchronous Boolean models. However, the problem is harder for the asynchronous updating scheme (see Section 2.2.3). Recently, Zañudo and Albert (2013) introduced a novel method to compute most asynchronous attractors.

Several methods have been proposed to specifically compute the stable states, for example, using constraint programing (Devloo et al., 2003) or polynomial algebra (Veliz-Cuba et al., 2010). To identify all the stable states, GINsim implements an efficient algorithm based on the manipulation of multivalued decision diagrams (see Naldi et al. (2007) for details). We will rely on this algorithm to compute the stable states of the T-helper cell differentiation model (see Section 3).

#### 2.2.3 Dynamical analysis – State Transition Graphs, representation and analysis

As mentioned above, the discrete dynamics of an LRG can be represented in terms of a *state transition graph* (STG), where the nodes denote *states* and the arcs represent *transitions* between states. A first approach to investigate dynamical properties consists in analyzing the STG in terms of attractors (terminal SCC), or regarding the existence of paths from an initial state towards specific attractors. The graph of strongly connected components (SCC) of the STG often provides a convenient, compressed view of the dynamics, in which attractors and reachability properties are easier to visualize. However, this representation may still encompass numerous single state components, hindering the interpretation of the dynamics. To further compress an STG and emphasize its topology, we recently proposed a novel representation, named *hierarchical transition graph* (see Bérenguier et al. (2013) for details).

Still, these representations do not address the identification of the attractors in large STGs. In this respect, Garg et al. (2008) proposed an efficient algorithm to identify all the attractors (synchronous and asynchronous schemes) of Boolean models. Their method relies on a binary decision diagram representation of the STG and can cope with very large models (Xie and Beerel, 1998). An implementation of this algorithm is available along with the software *genYsis* (http://www.vital-it.ch/software/genYsis/).

To further account for kinetic aspects, several strategies have been proposed. One strategy defines priority classes according to biologically founded time scale separations, *e.g.* fast versus slow processes (Fauré et al., 2006). Alternatively, time delays and constraints on them can be defined and handled with existing methods to analyze timed automata (Siebert and Bockmayr, 2006). Another approach consists in applying continuous time Markov processes on logical state spaces. Based on the delineation of a logical model along with a limited number of kinetic parameters, the software *MaBoSS* uses Monte-Carlo simulations to compute an estimate of the temporal evolution of probability distributions and of the stationary distributions of the logical states (Stoll et al., 2012). Finally, several authors proposed to consider differential models derived from logical models (Abou-Jaoudé et al., 2009; Mendoza and Xenarios, 2006; Wittmann et al., 2009).

### 2.3 Model checking for reachability analysis

#### 2.3.1 Model checking

The combinatorial explosion of the state spaces of discrete dynamical systems has been addressed during the last 30 years through the development of *model checking*, a computer science technique to verify properties in very large state spaces. The dynamics of discrete systems are directly mapped into a (graph-based) *Kripke structure* (Clarke et al., 1999). Model checkers receive a Kripke structure, either explicitly (representation equivalent to the STG), or implicitly in terms of a transition function specifying the successors of any given state. The latter case corresponds to *symbolic model checking*, which is handled by most model checkers nowadays. To perform a verification, a model checker takes as an input a set of properties denoting real-world observations, specified as temporal logic formulas, and verifies whether each of these properties is satisfied by the Kripke structure induced by the model under study.

Temporal logic formulas specify an order of sequences of transitions between states, without explicit time quantification. Several temporal logics have been defined with different expressive powers, using different types of operators. In the case of asynchronous updating, one might be interested in the study of each alternative path separately. This suggests the use of a temporal logic that provides path quantifiers where, at each step, a choice can be made between multiple paths, *i.e.*, a branching-time temporal logic. Within the family of branching-time temporal logics, *Computation Tree Logic* (CTL) is the most used one. Basic CTL operators are obtained by combining path quantifiers, Exists and All, with temporal operators, neXt, Future, Globally and Until (Clarke et al., 1999).

Different model checkers are available, differing in characteristics such as the underlying structure to represent the model dynamics or the supported temporal logics. A few examples are: CADP (Garavel et al., 2007), which uses labeled transition systems, supporting temporal logics with high expressive power like *Computation Tree Regular Logic* (CTRL) (Mateescu et al., 2011) or *μ*-calculus (Kozen, 1983); Antelope (Arellano et al., 2011), which uses state transition graphs, supporting Hybrid CTL, an extension of CTL with a special operator capable of selecting partly characterized states; and NuSMV (Cimatti et al., 2002), a symbolic model checker which uses multilevel decision diagrams, supporting the verification of properties through CTL or *Linear Temporal Logic* (LTL) (Clarke et al., 1999). As an open source project providing a generic description language for the specification of discrete dynamical systems, NuSMV is particularly prone to be extended by other research groups with additional features (see next subsection).

#### 2.3.2 Model checking applied to the analysis of logical models of signaling networks

Systems biology is a recent, successful application field for model checking techniques, covering a variety of modeling formalisms and/or type of properties to be verified (for details see Brim et al. (2013)). Here, we use GINsim, our modeling tool, which automatically exports biological models into NuSMV specifications. Biological observations are then expressed as sets of temporal logic formulas.

Computational models of signaling/regulatory networks aim at unraveling how external stimuli are processed to determine cell responses. In these networks, input nodes convey environmental cues, which are often assumed to be constant. Each combination of constant values of the inputs defines an STG, which is disconnected from the STGs defined by different combinations of input values. In other words, each fixed environmental condition defines a specific region of the state space in which the system is trapped. Rather than having input variables being part of the state definitions, we label each transition with the input values enabling this transition. This yields a state space defined solely by non-input variables and therefore a unique STG (Monteiro and Chaouiya, 2012). Extent of this reduction depends on the number of input components and on their value ranges.

In order to take advantage of this reduction, we need to be able to verify properties on both states and transition labels. NuSMV can only verify properties on state characterizations. We thus use *NuSMV-ARCTL*, which verifies properties combining both state and transition characterizations (Lomuscio et al., 2007). For the verification of such properties, *NuSMV-ARCTL* considers a CTL extension called *Action Restricted CTL* (ARCTL). Table 1 describes the syntax and semantics of the main ARCTL operators. With ARCTL, reachability properties are specified not only by characterizing the set of initial and target states, but also by constraining the values of some input components (transition labels), while remaining input components are allowed to freely vary.

**Table 1.** Syntax and semantics of the main ARCTL temporal operators (for a complete description see Lomuscio et al. (2007)); *α* denotes a restriction, defined only by the input variables, that must be satisfied (true) along the path; *ϕ* and *ψ* denote the restrictions, defined only by non-input variables, that must by satisfied at the target state or along the path.

| Syntax | Semantics |
| --- | --- |
| $EAF(\alpha)(\phi)$ | There is at least one path leading to a state that satisfies $\phi$ and the input restriction $\alpha$ must be satisfied along that path |
| $AAF(\alpha)(\phi)$ | All the paths lead to a state that satisfies $\phi$ and the input restriction $\alpha$ must be satisfied along all the paths |
| $EAG(\alpha)(\phi)$ | There is at least one path along which all the states satisfy $\phi$ and the input restriction $\alpha$ is satisfied along that path |
| $AAG(\alpha)(\phi)$ | All the states of all the paths satisfy $\phi$ and the input restriction $\alpha$ is satisfied along all the paths |
| $EA(\alpha)[\phi \text{ U } \psi]$ | There is at least one path along which all the states satisfy $\phi$, leading to a state that satisfies $\psi$ and the input restriction $\alpha$ is satisfied along the path |
| $AA(\alpha)[\phi \text{ U } \psi]$ | All the states of all the paths satisfy $\phi$, leading to a state that satisfies $\psi$ and the input restriction $\alpha$ is satisfied along all the paths |

Here, we take advantage of the expressiveness of ARCTL to study the influence of specific environmental conditions on the reprograming of chosen cell types (see Section 3). As presented hereafter, the T-helper cell differentiation model specified in GINsim is exported into a NuSMV specification, while properties of biological interest are specified as ARCTL temporal formulas. This allows us to define a novel, abstracted view of the dynamical behaviors called *reprograming graph*, which reveals switches between attractors upon changes in the input component values: the nodes of this graph represent the model attractors; and the arcs, labeled by specific combinations of input values, denote paths between those attractors.

## 3 Application: T-helper cell differentiation

T-helper (Th) (CD4+) lymphocytes play a key role in the regulation of the immune response. Upon activation by antigen presenting cell (APC), naive CD4 T cells (Th0) differentiate into specific Th subtypes producing different cytokines, which affect the activity of immune effector cell types (*e.g.* B lymphocytes, effector CD8 T cells, macrophages, etc.).

Three main types of signals are involved in this Th cell differentiation process (Supplementary Figure S1): (i) the presentation of antigenic peptide in conjunction with the major histocompatibility complex class II molecules (MHC-II) stimulate specific T cell receptors (TCR); (ii) co-stimulatory molecules further contribute to T cell activation and clonal proliferation; (iii) cytokines secreted by APCs and other cells bind their specific receptor(s) on the surface of Th0 cells, thereby affecting Th differentiation.

The cytokine environment instructs Th0 to enter a specific differentiation program in order to match the type of pathogen primarily stimulating the APCs. Over the last decade, a variety of Th subsets have been discovered (Nakayamada et al. (2012)), well beyond the initial identification of Th1 and Th2 dichotomy (Mosmann et al. (1986), Mosmann and Coffman (1989)).

Currently, several Th subsets (Th1, Th2, Th17, Treg, Tfh, Th9 and Th22) have been well established. These *canonical* subsets are characterized by their ability to express specific sets of cytokines under the control of a *master regulator* transcription factor (Supplementary Figure S1). However, various hybrid Th subsets expressing several master regulators have been recently identified (Ghoreschi et al. (2010), Duhen et al. (2012), Peine et al. (2013)). Evidences for substantial plasticity in Th differentiation have also been reported, including reprograming events between Th subsets under specific cytokine environments (Yang et al. (2008), Lee et al. (2009), Hegazy et al. (2010)). These findings challenge the classical linear view of Th differentiation and raise the question of which mechanisms underlie the observed diversity and plasticity of Th phenotypes.

Unraveling the complexity of Th differentiation and plasticity requires the development of an integrative and systematic approach articulating experimental analysis with computational modeling. We are currently setting a multi-parametric *in vitro* experimental approach to decipher how the microenvironment globally controls Th cell differentiation. In parallel, we are developing a comprehensive logical model of Th differentiation covering all parameters assessed in our experimental setup. Extending the modeling study reported in Naldi et al. (2010), the model presented here includes additional transcription factors and cytokine pathways and hence accounts for the differentiation of several novel Th subtypes. On the basis of this model, we illustrate how the computational methods described in Section 2, in particular model checking, can be used to assess biologically relevant dynamical properties. The model file as well as the steps to reproduce all the results described below are available from the model repository of the GINsim web site.

### 3.1 Model description

Our Th differentiation model encompasses different layers (see Figure 2), namely:

- the cytokine inputs along with the Antigen Presenting Cells (APC);
- the cytokine receptors and their subchains, along with the T cell receptor (TCR) and the costimulatory receptor CD28;
- the intracellular signaling factors, including ”Stat” family proteins (Stat1, Stat3, Stat4, Stat5, Stat6), the TCR and co-stimulatory signaling components (NFAT, I*κ*B, NF*κ*B), the master regulators (Tbet, Gata3, Ror*γ*t, Foxp3, Bcl6), along with additional transcription factors involved in Th differentiation (cMaf, PU.1, Smad3, IRF1, Runx3);
- the main cytokines secreted by Th cells;
- a component modeling the proliferation of the cell.

By and large, the model encompasses 21 signaling pathways (comprising external cytokines, receptor chains, etc.), 17 transcription factors, 17 cytokines expressed by Th cells, and one node accounting for cell proliferation, amounting to 101 components in total. In comparison with the model reported in (Naldi et al., 2010), this model integrates factors characterizing novel Th subsets (Tfh, Th9 and Th22) as well as additional signaling pathways and secreted cytokines involved in the differentiation and the definition of Th cellular types. A complete list of the components of the model along with supporting evidence is provided in Supplementary Table S1. The logical rules associated with the components are listed in the Supplementary Table S2.

As in Naldi et al. (2010), a gene expression pattern is associated with each canonical Th subtype, based on experimental evidence (Table 2). Each pattern represents a restriction of Th cell states to a subset defined by the activation or the inactivation of critical markers characterizing the corresponding canonical Th subtype. In the following sections, we present the results obtained by the application of the aforementioned computational methods to our Th differentiation model.

**Table 2.**
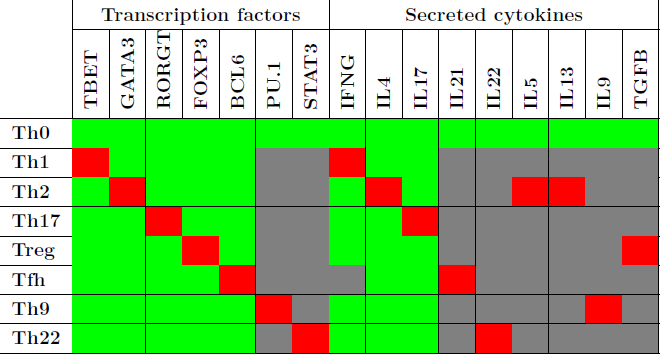
Logical expression patterns for the canonical Th subtypes. Red and green cell denote the activation and inactivation of the components (column entries), with respect to the canonical Th subtype (row entries). Gray cells represent components that can be either activated or inactivated for the corresponding canonical Th subtype. The components not mentioned are considered to be either activated or inactivated, except in the case of Th0, where they are all inactivated.

### 3.2 Static analysis

We first checked the consistency of the rules inferred from experimental data (Supplementary Table S2) with the interactions composing the regulatory graph of Figure 2. An analysis of interaction functionality led to the identification of a single non-functional interaction (IL10R → Stat3). Although the role of this interaction is not yet clear, we kept it in the regulatory graph as it is documented (see Supplementary Table S1).

Next, to ease the model analysis, we derived a reduced version of this model using the reduction described in Naldi et al. (2011), keeping internal components characterizing the canonical Th patterns (cf. Figure 2, where the grey nodes denote the components selected for reduction).

Using the method described in Naldi et al. (2007), we computed all the stable states for all the input combinations and grouped them according to phenotypic markers (see also subsection 2.2.2 above). Since the reduction preserves the stable states, each stable state of the reduced model strictly corresponds to one stable state of the original model (and vice versa). This analysis led to the identification of 82 context-dependent stable states, including sets of stable states matching the activity patterns associated with each canonical Th type (see Supplementary Table S3). This analysis further predicts the existence of stable states representing hybrid cellular types, *i.e.* expressing several master regulators, including four hybrids expressing two master regulators, which have been recently reported in the literature, and another one (Tbet^+^Gata3^+^Foxp3^+^) expressing three master regulators, which has not yet been experimentally observed. Each of the stable states found is associated with a subset of input combinations. One can actually recover the input configurations associated with each stable state, getting a first insight into the role of environmental cues in controlling the asymptotic behaviors of the system (see Section 3.3.2 for an illustration of this analysis).

### 3.3 Reachability analysis

As mentioned above, static analysis of the logical model allows for the identification of stable Th cellular types along with their associated input configurations. Our next aim is to determine how environmental cues control the differentiation and plasticity of these Th cell types. This question amounts to check whether a cellular type is reachable from a given initial state for specific input conditions. This kind of questions can be efficiently addressed using model checking, by verifying temporal properties under constant or varying input conditions.

We first carried out a systematic analysis of reachability properties between the canonical Th subsets as defined in Table 2, under specific constant polarizing cytokine environments. We consider nine prototypic environmental conditions (listed in Table 3) for this reachability analysis, including seven documented polarizing cytokine environments known to commit Th cells into the canonical subsets.

**Table 3.**
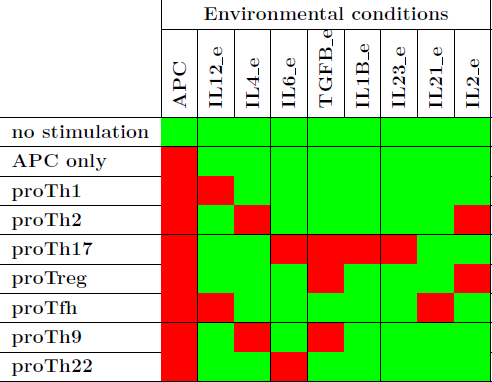
Prototypic environmental conditions. Each row corresponds to a prototypic environment defined as a combination of APC and cytokine inputs (columns). These environments encompass seven documented polarizing environments (denoted “proThX”) known to polarize naive Th cells into the canonical subsets (defined in Table 2). Red/green cells represent present/absent inputs. Non mentioned inputs are considered as absent.

We used the *NuSMV-ARCTL* model checker and instantiated the following generic property with values from Table 2 and Table 3:

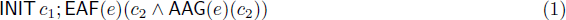

This property asserts the existence of a path from a canonical Th pattern *c*_1_, instantiated with values from Table 2, towards a (stable) canonical Th pattern *c*_2_, also instantiated with values from Table 2, under an input condition *e*, instantiated with values from Table 3.

Checking this property for all the combinations of canonical Th patterns and input conditions, one can represent the verified properties through a reprograming graph, which here abstract paths between Th patterns and recapitulate the polarizing and reprograming events predicted by our model (Figure 3). This graph provides a global and synthetic representation of Th plasticity depending on environmental cues. Focusing on polarizing events from naive Th0 cells to the other Th subtypes, our model is consistent with experimental data, showing that each canonical subset can be reached from the naive state Th0 (blue arcs starting from Th0 in Figure 3) in the presence of specific polarizing cytokine combinations (denoted by the labels associated with the blue arcs in Figure 3). The remaining Th-subtype conversions present in the reprograming graph would need to be assessed experimentally.

**Figure 3.**
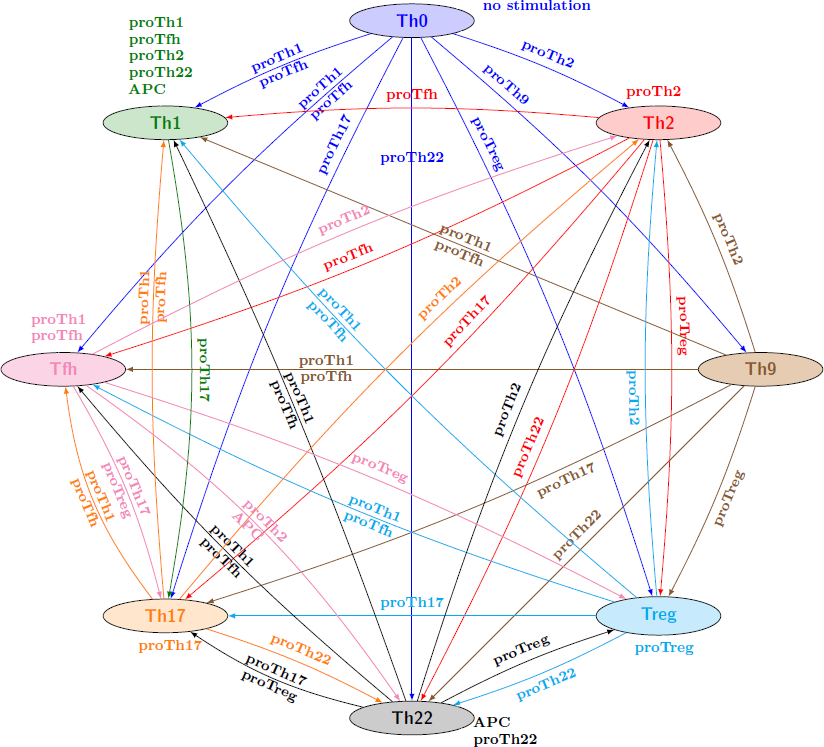
Reprograming graph, considering all canonical Th subtypes, generated with the model checker *NuSMV-ARCTL.* Nodes represent sets of states characterizing the canonical Th subtypes defined in Table 2. There is an arc labeled with *e*, going from node *c*_1_ to node *c*_2_, whenever the following ARCTL temporal logic formula is verified: INIT *c*_1_; EAF(*e*)(*c*_2_ ∧ AAG(*e*)(*c*_2_)). It should be noted that the existence of a single reprograming path from a Th subtype to another one does not necessarily imply the stability of the target Th subtype, since NuSMV-ARCTL considers that a property is true if and only if it is verified by the whole set of states in the initial conditions. Hence, if at least one state associated with a given subtype points to a state not associated with this subtype (for given input conditions), then the stability of the Th subtype is not represented (see for example Th9 subtype, which is not considered stable under proTh9 input condition).

An extensive discussion of all these Th type conversions is beyond the scope of this article. However, one interesting outcome is the inherent dissymmetry of this graph, with some Th subtypes apparently very stable under the environments considered (*e.g.*, Th1 node, with seven incoming arcs but only one outgoing one), while others need very specific conditions for their maintenance (*e.g.*, Th9 node, with six outgoing arcs and only one incoming one).

Hereafter, we focus on specific biological questions regarding Th differentiation and plasticity and show how model checkers can be applied to address these questions. Two biological questions will be considered: (i) the delineation of reprograming strategies to convert Th1 into Th2, and vice versa; (ii) the identification of relevant environmental conditions enabling the polarization to the Tbet^+^Gata3^+^Foxp3^+^ hybrid Th subtype identified in the course of the stable state analysis.

#### 3.3.1 Reprograming between Th1 and Th2

Since the discovery of Th1 and Th2 subsets, Th1 and Th2 commitments have been for a long time considered as mutually exclusive (Murphy and Reiner, 2002). However, recent experimental observations challenged this Th1/Th2 dichotomy (Antebi et al., 2013; Hegazy et al., 2010; Peine et al., 2013), raising the question of which environmental conditions can instruct Th1 or Th2 interconversions.

We first address this question by investigating Th1-Th2 reprograming strategies for the prototypic input conditions (listed in Table 3). From the reprograming graph (Figure 3), two strategies emerge: (1) although there is no direct path from Th1 cells towards Th2 cells, one could consider a two-step approach to reprogram Th1 cells into Th2 cells by applying a proTh17 condition, followed by a proTh2 condition; (2) as there is a direct path from Th2 to Th1 labeled with proTfh conditions, the application of a proTfh environment would potentially reprogram Th2 cells into Th1 cells.

We then ask whether other (constant or varying) input condition strategies could be identified for the reprograming between Th1 and Th2, beyond the prototypic environmental conditions. This question can be addressed using the following ARCTL formulas:

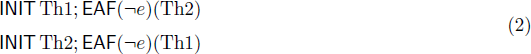

where *e* denotes the set of all the prototypic inputs (and consequently ¬*e* denotes the set of all the input combinations except the prototypic ones). *NuSMV-ARCTL* evaluates both formulas as true, implying that it must exist at least one non-prototypic (constant or varying) input condition allowing for the reprograming of Th1 into Th2, and vice versa.

To further illustrate the power of model checking to analyze cell plasticity, we focus on Th2 reprograming into Th1. Our initial analysis predicts that the prototypic proTh1 cytokine environment does not enable this reprograming (see Figure 3). However, looking more closely at the regulatory graph, we see that TGF*β* signaling pathway inhibits Gata3, the master regulator of Th2 cells (Figure 2). This suggests an alternative, two-step strategy, to reprogram Th2 into Th1, by applying first TGF*β* in the cell environment to inhibit Gata3, and thereby block its inhibitory effects on Th1 differentiation, followed by the application of a proTh1 environment to induce Th1 polarization. We can assess this strategy using the following ARCTL formula:

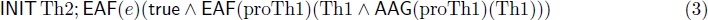

where *e* is an input condition restricting only TGF*β* to ON (all other inputs can freely vary). This property is evaluated as true. We can thus conclude that this alternative strategy could also be used to reprogram Th2 into Th1 cells.

Beyond this analysis, one can further investigate network perturbations (*e.g.* gene knock-in or knockout) enabling Th1-Th2 reprograming. This type of question can be assessed using model checking of perturbed models. Here, we focus again on reprograming Th2 cells into Th1 cells under the prototypic proTh1 input condition. Over-expression of a Gata3 (Th2 signature) inhibitor (*e.g.*, PU.1 or Bcl6) would be a relevant option. However, Bcl6 should be discarded because it also inhibits Tbet (Th1 signature) (cf. the logical rule of Tbet in Supplementary Table S2). Using the generic property (1), the analysis of a perturbed model with ectopically expressed PU.1 suggests that this perturbation can indeed induce the reprograming of Th2 into Th1 in the presence of the prototypic proTh1 input condition.

Finally, we can study the role of critical regulatory interactions underlying such reprograming events through model checking analyses of perturbed models. Turning back to the reprograming strategies 1 and 2 presented above, we now focus on the inhibitory interactions acting upon Tbet and Gata3, the master regulators of Th1 and Th2 cell types, respectively. For example, in Figure 2, we see that Ror*γ*t inhibits Tbet, which could be relevant for reprograming strategy 1, while Bcl6 inhibits Gata3, which might be relevant for reprograming strategy 2. Analyses of perturbed models, using the ARCTL generic property (1), where either one or the other interaction is suppressed, suggest that the inhibition of Tbet by Ror*γ*t is indeed necessary for reprograming strategy 1, whereas the inhibition of Gata3 by Bcl6 is indeed necessary for reprograming strategy 2.

#### 3.3.2 Reachability of the triple hybrid subtype Tbet^+^Gata3^+^Foxp3^+^

The steady state analysis of our model in Section 3.2 predicts the existence of a stable hybrid Th subtype co-expressing Tbet (characteristic of the Th1 signature), Gata3 (Th2 signature) and Foxp3 (Treg signature), which has not been yet experimentally reported.

Using model checking, we can evaluate environmental conditions that might enable the polarization of naive Th0 cells into this hybrid subtype. First, the input combinations for which this hybrid subtype is stable can be extracted directly from the steady state analysis (not shown). In these combinations, some cytokines appear to be either always ON, namely IL15, or always OFF, TGF*β*. Moreover, TGF*β* signaling, via Smad3, is clearly needed to activate Foxp3 (see logical rule of Foxp3 in Supplementary Table S2), suggesting that a transient TGF*β* environment is necessary to polarize naive Th0 cells into the hybrid subtype Tbet^+^Gata3^+^Foxp3^+^. This last hypothesis can be verified using the ARCTL formula:

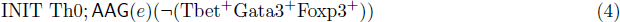

where *e* denotes an input condition restricting only *β* to OFF (all other inputs can freely vary). This formula states that the hybrid pattern cannot be reached from whatever path leaving the canonical Th0 pattern under the input restriction *e*. As the property is evaluated as true, we conclude that a strategy without (transient) *β* in the environment cannot polarize Th0 into the hybrid subtype, confirming our hypothesis.

Therefore, a two-step approach to polarize naive Th0 cells into the hybrid subtype Tbet^+^Gata3^+^Foxp3^+^ could be considered, applying *β* transiently, before applying an environment containing IL15. This strategy can be evaluated using the ARCTL formula:

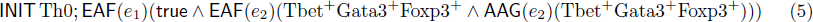

where *e*_1_ denotes the first input combination (in which TGF*β* and APC are ON), and *e*_2_ denotes the second input combination (in which IL15 and APC are ON and TGF*β* is OFF). Two additional input cytokines were also considered in these combinations: IFN_*γ*_ for Tbet activation and IL25 for Gata3 activation. We consider 18 strategies (input configurations), including six of them being able to polarize Th0 into the hybrid subtype (see Supplementary Table S4). Interestingly, these six strategies have all IFN_*γ*_ switched OFF in the first input combination and turned ON in the second input combination.

## 4 Conclusions and prospects

Considering logical models of large cellular regulatory networks, we have focused on model checking to explore induced dynamical properties. Over the last decades, computer scientists have made spectacular advances in the development of powerful model checkers, regarding both performances and expressivity power. Several model checkers are freely available and can be used to check specific properties of dynamical models of biological systems. As illustrated above, asynchronous dynamics of logical models integrating signaling pathways with transcriptional networks can be readily translated into explicit or implicit Kripke structures, and thereby become amenable to standard or action-restricted model checking.

We have applied this approach to the analysis of a logical model for a comprehensive signaling/regulatory network controlling T-helper (Th) cell differentiation, which encompasses 101 components (most but not all Boolean) and 221 regulatory interactions. As the state space induced by this network is gigantic (encompassing over 2^100^ states), scalable formal methods enabling the exploration of interesting dynamical properties are paramount. In this respect, we have combined three complementary approaches: (i) a formal reduction method conserving the main dynamical properties, including the stable states (described in Section 2.1.3); (ii) an algorithm enabling the identification of all the stable states in large logical models (described in Section 2.2.2); (iii) the use of model checking to verify the reachability of specific stable patterns (reprograming of specific Th cell subtypes) from given initial conditions, in the presence (or not) of network perturbations.

We have illustrated the power of the model checking approach by addressing key biological questions related to Th differentiation and plasticity in response to environmental cues. To this end, we have formulated two main types of queries: (i) is it possible to reprogram a specific Th subtype into another one, using specific fixed (or any free) cytokine combinations, in a single (or a multiple) step(s)? (ii) does such reprograming depend on specific regulatory components or interactions (using perturbed models)? We have shown that such biological questions can be efficiently assessed using action-restricted model checking. Using the model checker *NuSMV-ARCTL*, we could confirm that our model is consistent with the polarization of naive Th cells into the canonical Th subsets under specific cytokine input environments, and delineated several strategies allowing the reprograming between specific Th subtypes (Th1 and Th2) as well as the polarization of naive Th cells towards a novel Th hybrid subtype predicted by our analysis (Tbet^+^Gata3^+^Foxp3^+^).

Although our logical model for T-helper cell differentiation should be further refined using a comprehensive experimental data set (work in progress), it could be already used as a framework to design informative experiments regarding the identification of Th hybrid subtypes, or yet to characterize Th cell plasticity. Some of the resulting predictions (*e.g.* the existence of Tbet^+^Gata3^+^Foxp3^+^ Th hybrid) currently serve as a basis to design experiments *in vitro.*

More generally, we wish to stress that formal modeling can be used at various stages of the deciphering of complex regulatory networks, provided that the formal framework and methods used, as well as the modeling scope, are adapted to the data available. In this respect, qualitative (Boolean or multivalued) logical modeling is well suited to model large biological regulatory networks, for which reliable quantitative data are often lacking (Saez-Rodriguez et al. (2007), Grieco et al. (2013)).

Beyond the proof of concept, the development of user-friendly tools is required for a wider use of model checking in systems biology. In this respect, we are currently working on improving the interaction between GINsim and NuSMV-ARCTL in two distinct ways, which will be made available in a forthcoming release of GINsim: (1) implementing recurrent temporal logic patterns into our software GINsim to ease the definition of temporal logic formulas; (2) automating the interaction with the model checker and the parsing of the results, as well as the generation of the reprograming graph.

## Author Contributions

The model was defined by WAJ with the help of MG under the supervision of VS and DT. All model analyses were performed by WAJ and PTM under the supervision of CC and DT. AN wrote some of the scripts and implemented several algorithms used in this study. All authors contributed to the writing of the manuscript and agreed with its final content.

## Acknowledgement

WAJ has been supported by postdoctoral grants from the LabEx MemoLife and from the Ecole Normale Supérieure. PTM was supported by the FCT grants PEst-OE/EEI/LA0021/2013 and IF/01333/2013. MG was funded by the PhD program of the Agence National de Recherche sur Le Sida (ANRS).

